# Dissociation of Striosome and Matrix Activation in the Human Striatum During the Cue and Execution Phases of Working Memory

**DOI:** 10.64898/2026.01.29.702581

**Authors:** Alishba Sadiq, Jeff L. Waugh

**Affiliations:** Division of Pediatric Neurology, University of Texas Southwestern, Dallas, TX, USA; Department of Pediatrics, and Department of Neurology, University of Texas Southwestern, Dallas, TX, USA

**Keywords:** Striatum, Compartment, Striosome, Matrix, Connectivity-based parcellation, Task-based fMRI, Working memory

## Abstract

The striatum comprises two neurochemically and anatomically distinct tissue compartments, the striosome and matrix, that are hypothesized to support different aspects of cognition and action. In animal studies, the striosome has been linked to reward evaluation, emotional learning, and decision-making under conflict, whereas the matrix is more closely associated with sensorimotor integration and task execution. However, evidence for compartment-specific function in humans is limited and indirect. Using probabilistic tractography, we identified voxels with striosome-like and matrix-like patterns of structural connectivity in healthy adults. We then examined how these compartment-like voxels responded to task demands during an fMRI *n*-back working-memory paradigm that visually presented four stimulus categories (*body* part, *face, place*, or *tool*). We assessed activation in a low-load condition (0-back, remembering a just-viewed stimulus) vs. a high-load condition (2-back, remembering a stimulus viewed two prior). Functional activation was temporally segregated and matched our prior findings in motor tasks: striosome-like voxels were preferentially engaged during the cue and initial preparation phases, whereas matrix-like voxels dominated during task execution. Trial accuracy strongly modulated striatal activation, with both compartments showing significantly greater responses during “correct” than in “error” trials. Notably, the accuracy-related increase in activation was larger in striosome-like voxels, consistent with a prominent role for striosomal processing in performance evaluation. Both striosome- and matrix-like voxels significantly increased activation from 0-back to 2-back, indicating sensitivity to working-memory load, with larger increases for matrix-like than for striosome-like voxels. Category-selective responses also differed by compartment and cognitive load. Under low working-memory load (0-back), stimulus-category effects were modest and broadly similar between compartments. Under higher load (2-back), activation in striosome-like voxels remained selective for specific stimulus categories, while matrix-like voxels lost category specificity. Together, these findings suggest that the striosome–matrix distinction generalizes from motor to cognitive domains, reflecting a conserved division between preparatory and execution-related processes that varies systematically with task demands, memory category, and performance accuracy. This convergence of compartment-specific responses across domains points to a core organizational principle of the human striatum with potential implications for neuropsychiatric diseases.

**Key Points:**

- Striatal medium spiny neurons are organized into two interdigitated compartments, the striosome and matrix, which are embryologically, pharmacologically, and anatomically distinct. Compartment-specific functions have been demonstrated in animals, but their roles in human cognition are unexplored.
- We found that in humans, striosome-like voxels preferentially activated during memory cues, while matrix-like voxels preferentially activated during recall and memory maintenance. In both compartments, activation scaled with task difficulty.
- Activation in striosome-like voxels scaled more strongly with task accuracy and difficulty, suggesting a striosome-selective role in vigilance and/or motivation.

## 1 Introduction

Historically, the basal ganglia have been primarily associated with motor control, a view shaped by early 20th-century work by Kinnier Wilson (Wilson 1912, Wilson 1914) and Vogt (Vogt 1911), who observed motor impairments following damage to this region. The cardinal motor symptoms of Parkinson’s disease (PD) arise from the progressive degeneration of dopaminergic neurons in the substantia nigra pars compacta (SNc), leading to dopamine (DA) depletion in the striatum (Surmeier, Graves and Shen 2014). However, anatomical and functional evidence accumulated over the past several decades has established that the striatum supports not only motor control but also cognitive and limbic functions. As the major input nucleus of the basal ganglia, the striatum is thought to mediate the selection and sequencing of motor actions (Mink 1996, Graybiel 2008), it evaluates cortical “action plans,” integrates sensory and motivational context with prior experience, and transmits this evaluation to downstream nuclei that shape motor output through temporally patterned inhibitory signaling to the thalamus and brainstem. Anatomical studies show that the striatum forms part of several cortico– striato–thalamo–cortical circuits, each responsible for processing different types of information such as motor, cognitive, or limbic functions (Middleton and Strick 2000). Haber et al. (2000) further demonstrated in macaques that the ventral midbrain acts as an interface, enabling information exchange between these striatal regions and integrating signals related to movement, cognition, and emotion (Haber, Fudge and McFarland 2000). These findings highlight the role of the striatal regions to cognitive as well as motor functions.

Basal ganglia dysfunction contributes not only to classic movement disorders but also to a range of cognitive and psychiatric impairments. For example, in Parkinson’s disease and Huntington’s disease (Wichmann and Dostrovsky 2011) motor symptoms are typically accompanied by, and are often preceded by, significant cognitive and mood symptoms. In addition, conditions such as obsessive–compulsive disorder (OCD; (Burguiere, Monteiro et al. 2015)), attention-deficit/hyperactivity disorder (ADHD; (Singh, Skippen et al. 2024)), Tourette’s syndrome (TS;(Albin 2006)), and depression (Dunlop and Nemeroff 2007) highlight the basal ganglia’s central involvement in cognitive control and behavioral regulation. Within this system, the striatum (comprised of caudate and putamen) integrates afferent signals from the cortex, thalamus, and brainstem to guide the selection and execution of discrete behaviors. Striatal spiny projection neurons (SPNs) are organized into two distinct tissue compartments, the striosome and matrix, which differ in their developmental origins, neurochemical markers, pharmacological properties, and connectivity with other brain regions (Crittenden and Graybiel 2011). The striosome is a labyrinthine, 3-dimensional structure that is surrounded by the matrix (Brimblecombe and Cragg 2017). The compartments are interdigitated and the precise location of the striosome branches varies between individuals, precluding the use of region-of-interest approaches to distinguish the functions of striosome and matrix.

The striosome receives dense inputs from limbic and prefrontal regions and exerts direct inhibitory control over dopaminergic neurons in the substantia nigra pars compacta, positioning the striosome to regulate motivational salience and the gating of task-relevant information (Watabe-Uchida, Zhu et al. 2012, Friedman, Homma et al. 2015, Crittenden, Tillberg et al. 2016). In contrast, the matrix receives inputs from dorsolateral prefrontal, parietal association, and sensorimotor areas, and projects to basal ganglia output nuclei, supporting the sustained maintenance and manipulation of information necessary for cognitive performance (Lévesque and Parent 2005, Watabe-Uchida, Zhu et al. 2012). Dopaminergic inputs further differentiate the compartments, decreasing striosome firing while increasing matrix firing (Prager et al., 2020). Together, this architecture suggests that striosome and matrix, and the functional networks in which they are embedded (Sadiq, Funk and Waugh 2025), form complementary but distinct substrates for higher-order cognitive functions, including working memory (Weglage, Wärnberg et al. 2021).

Despite detailed descriptions of this two-compartment architecture in animal models and human histology, its functional significance for memory has yet to be elucidated. Evidence from animal and human studies supports functional specialization between the striosome and matrix compartments. The striosome directly regulates nigral dopamine signaling and mediates decision-making under conflict (White and Hiroi 1998, Friedman, Homma et al. 2015), aversive learning, and negative-valence memory (Jenrette, Logue and Horner 2019, Nadel, Pawelko et al. 2020). In contrast, the matrix receives extensive sensorimotor and associative inputs and is critical for movement execution, habit formation, and behavioral routines (Aosaki, Graybiel and Kimura 1994, Lévesque and Parent 2005). Through its strong projections to both the direct and indirect pathways, the matrix compartment contributes to an oppositional mechanism in which one pathway facilitates desired motor programs (the direct pathway) while the other suppresses competing or inappropriate ones (the indirect pathway), thereby enabling precise action selection and control (Mink 1996). Importantly, this dynamic has also been proposed to extend to cognitive domains, including attention, working memory, and rule selection (Swerdlow and Koob 1987, Graybiel 1997, Bejjani, Damier et al. 1999). Within the context of working memory, such circuitry would allow the matrix to sustain task-relevant representations through the direct pathway while concurrently inhibiting competing representations via the indirect pathway. This balance between stabilization and suppression provides the neural substrate for selective flexibility—the ability to maintain focus on current goals while efficiently shifting to alternative strategies when required.

We recently demonstrated, using probabilistic diffusion tractography, that connectivity-based parcellation can delineate striatal voxels with striosome-like or matrix-like properties in vivo, based on their distinct cortical and subcortical connectivity profiles (Waugh, Hassan et al. 2022). This approach enables individualized mapping of compartment-like organization in the living human brain, while recognizing that “striosome-like” and “matrix-like” voxels are defined probabilistically and are not the equivalent of compartment mapping by immunohistochemical markers. Nevertheless, MRI-based parcellations recapitulate all of the anatomic features of striosome and matrix architecture described in tissue. First, the relative abundance of matrix-like (∼85%) and striosome-like (∼15%) voxels closely matches histological estimates from human and primate studies (Desban, Kemel et al. 1993, Holt, Graybiel and Saper 1997, Waugh, Hassan et al. 2022, Funk, Hassan et al. 2023, Funk, Hassan and Waugh 2024, Sadiq, Funk and Waugh 2025, Waugh and Tieu 2025). Second, their spatial distributions mirror histological observations, with striosome-like voxels concentrated in the rostroventral striatum and matrix-like voxels more evenly distributed in dorsolateral and caudal regions (Graybiel and Ragsdale Jr 1978). Third, connectivity of these voxels is organized somatotopically (Funk, Hassan et al. 2023, Sadiq, Funk and Waugh 2025, Waugh and Tieu 2025), consistent with tract tracing studies in animals (Flaherty and Graybiel 1993, Eblen and Graybiel 1995). Fourth, matrix-like voxels tend to form large contiguous clusters, whereas striosome-like voxels are more often isolated (Waugh and Tieu 2025). Finally, this parcellation method is highly reliable, with a test– retest error rate of only 0.14% across repeated scans (Waugh, Hassan et al. 2022). Importantly, both structural and functional analyses have shown that these compartment-like biases depend on the precise spatial location of voxels rather than their striatal “neighborhood”: even small (2-3 mm) random displacements abolish compartment-specific structural connectivity patterns (Funk, Hassan et al. 2023, Sadiq, Funk and Waugh 2025) and resting state functional connectivity networks (Sadiq and Waugh 2025). Together, these findings establish a reproducible framework for probing the functional significance of striosome- and matrix-like voxels in humans, including their roles in working memory.

Although striosome and matrix have been well described in animal studies and human histology, their contributions to higher-order cognition remain poorly understood. It is unclear how the compartments participate in the dynamic processes of working memory, such as preparing to update task-relevant information, sustaining representations across delays, and executing responses based on those representations. We previously demonstrated that striosome-like and matrix-like voxels are embedded in segregated structural and resting-state functional networks (Funk, Hassan et al. 2023, Funk, Hassan and Waugh 2024, Sadiq, Funk and Waugh 2025). These findings suggest that the two compartments may form distinct large-scale systems, consistent with hypotheses that striosome dysfunction contributes to neuropsychiatric disorders through its unique limbic and dopaminergic connectivity (Crittenden and Graybiel 2011). Such organization provides a plausible circuit basis for hypotheses that striosome dysfunction contributes to neuropsychiatric disorders, particularly those involving impaired motivation and valuation. However, whether striosome- and matrix-like voxels are differentially recruited during working memory tasks in humans has not been established. To address this gap, we used compartment-like voxels as seeds in task-based fMRI, testing whether the two compartments exhibit distinct activation profiles during working memory tasks. Specifically, we asked: (1) Are striosome- and matrix-like voxels differentially engaged during working memory demands? (2) Do these differences depend on distinct phases of the task, such as cue-related processing versus information maintenance and response execution? (3) Do the observed dynamics align with proposed roles of the striosome in motivational and evaluative processing, and of the matrix in sustaining task-relevant cognitive representations?

This study provides the first direct evidence of functional dissociation between striosome- and matrix-like activation in the human striatum during working memory, with activation differences that scaled with working-memory demands, offering a new framework for linking microcircuit-specific striatal activation patterns to higher-order cognition and neuropsychiatric symptoms.

## 2 Materials and Methods

Overview: we combined structural and functional connectivity approaches to investigate compartment-specific activation in the human striatum during working memory tasks (Figure 1). We perfomed structural connectivity-based parcellation using probabilistic diffusion tractography to identify striatal voxels with striosome-like or matrix-like connectivity biases. To assess functional activation, we extracted task-evoked BOLD signals from each subject’s striosome- and matrix-like masks. Each participant’s response to working memory events was modeled using a general linear model (GLM), a standard framework for estimating task-related brain activity. We calculated contrast parameter estimates (COPEs) at each voxel to quantify activation differences between task and baseline conditions, yielding compartment-specific activation measures. This framework allowed us to compare how the magnitude and timing of activation differed between striosome- and matrix-like voxels across varying working-memory demands.

**FIGURE 1:**
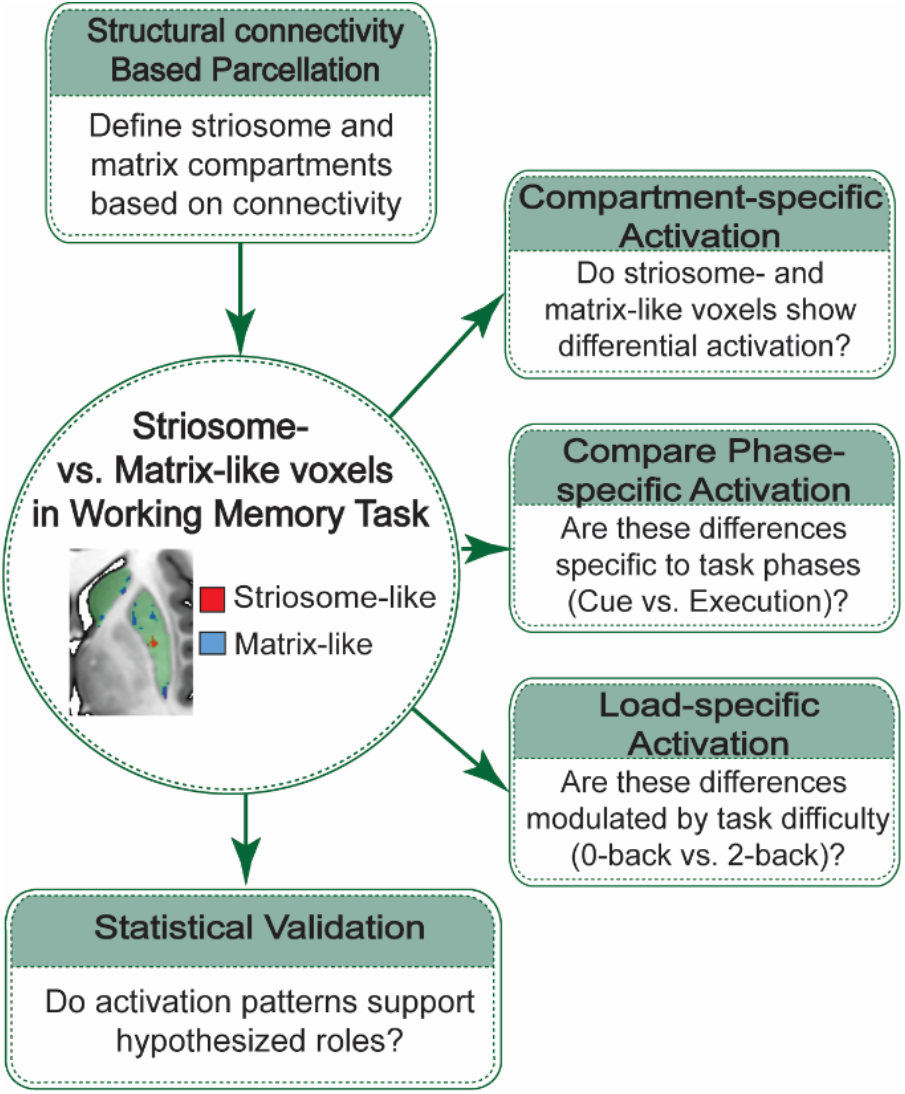
Schematic of the experimental workflow for evaluating compartment- and load-specific activation in the human striatum. Individual striosome-like and matrix-like compartments were delineated using structural connectivity–based parcellation. Because precise striosome location varies between individuals, uniform region-of-interest approaches cannot reliably distinguish the compartments. Therefore, subject-specific compartment-like masks were used to extract activation during a working-memory task.

### 2.1 Study Population

This study was a secondary analysis of MRI data from the Human Connectome Project (HCP) S1200 release (Van Essen, Smith et al. 2013), which includes high-quality multimodal neuroimaging data from a large cohort of healthy young adults. From the original sample of 1,206 participants, we included only those with complete diffusion MRI and task-based (working memory) fMRI data. Participants were excluded if they reported any lifetime history of illicit or addictive substance use (including cocaine, hallucinogens, cannabis, nicotine, opiates, sedatives, or stimulants). We also excluded individuals who met DSM-5 criteria for Alcohol Use Disorder (Alcohol Abuse or Dependence) or who reported consuming, on average, more than four alcoholic drinks per week in the year prior to scanning. All participants provided written informed consent as part of their enrollment in the HCP study (Van Essen, Ugurbil et al. 2012). In a related study using an overlapping experimental cohort, we previously identified resting-state networks that covaried with striosome-like versus matrix-like voxels (Sadiq, Funk and Waugh 2025).

### 2.2 MRI Acquisition Protocols

Task-based functional MRI (tfMRI) and diffusion tensor imaging (DTI) scans were obtained from the Human Connectome Project (HCP) S1200 dataset. All were collected on 3T MRI systems using standardized protocols across three participating sites. The tfMRI scans were acquired with echo-planar imaging (EPI) sequences matched to the resting-state fMRI (rs-fMRI) acquisitions, ensuring identical spatial and temporal resolution between modalities. A multiband gradient-echo EPI sequence was employed with the following acquisition parameters: repetition time (TR) of 720 ms, echo time (TE) of 33.1 ms, flip angle of 52°, field of view (FOV) of 208 × 180 mm, and matrix dimensions of 104 × 90, yielding 72 contiguous axial slices. The multiband acceleration factor was 8, with an echo spacing of 0.58 ms and a bandwidth of 2290 Hz per pixel. The tfMRI resolution was 2.0 mm isotropic, which we have previously demonstrated is sufficient to resolve striosome-like and matrix-like structural connectivity (Waugh, Hassan et al. 2022). Each working memory task condition included two runs of 405 frames, each lasting approximately 5 minutes and 1 second. The short scan duration and fast TR enabled high temporal resolution for task-evoked activity mapping. DTI data for S1200 subjects was acquired at 1.25 mm isotropic resolution using 200 directions (14 B0 volumes, 186 volumes at noncolinear directions) with the following parameters: repetition time = 3.23 s, echo time = 0.0892. DTI scans included both anterior-posterior and posterior-anterior acquisitions, allowing for correction of susceptibility artifacts.

#### 2.2.1 Working Memory-Task Design

The experiment used a working-memory task that incorporated stimuli from multiple visually-presented categories (e.g., *faces, places, body* parts, *tools*) under both 0-back and 2-back conditions. In the 0BK task, participants identified a pre-specified target, whereas in the 2BK task they determined whether the current stimulus matched the one presented two trials earlier. Stimuli consisted of pictures of *places* (e.g., outdoor scenes or buildings), *tools, faces*, and non-mutilated *body* parts (with no nudity), presented in separate blocks within each run. Each run included eight task blocks: half using the 0-back and half the 2-back task, and four fixation blocks. At the start of each block, a 2.5-second cue indicated the task type (and the target for 0-back). Task blocks contained 10 trials (each 2.5 seconds: 2 seconds stimulus plus 500 ms inter-task interval), totaling 25 seconds per block, while fixation blocks lasted 15 seconds. The fixation blocks served as baseline condition, during which participants fixed on a centrally presented cross without performing any task. All memory task events were modeled relative to the average of the four 15-second fixation blocks. This approach allowed us to contrast memory-related activation with a stable, aggregated baseline estimate.

For subsequent analyses, we examined activation patterns as a function of both task performance and compartment type. For the accuracy-based analysis, activation values were extracted separately for correct and error trials within each compartment and aggregated across stimulus categories. Not all participants contributed data to every condition: all 965 participants had valid 0-back correct trials, whereas only 844 had sufficient 2-back error trials to estimate activation reliably. Because 0-back error trials were rare, we restricted our assessment of error-related activation to the 2-back condition. Category-selectivity analyses contrasted activation within each compartment for each stimulus category (e.g., *place*) against the mean of the other three categories (e.g., *tools, faces*, and *body* parts).

### 2.3 fMRI Preprocessing

We analyzed the minimally preprocessed HCP motor task-fMRI data (Glasser, Sotiropoulos et al. 2013). The HCP minimal preprocessing pipeline applies gradient-distortion correction, motion correction (rigid-body realignment), EPI distortion correction using spin–echo field maps, and registration of each subject’s functional data to their T1-weighted anatomical image and subsequently to MNI space via the fMRIVolume pipeline. The data is further cleaned using ICA-FIX, which removes structured noise components, including motion-related artifacts. No additional preprocessing steps, such as temporal filtering or spatial smoothing were applied before first-level modeling.

To evaluate residual motion, we computed framewise displacement (FD) following Power et al. (2012) using the HCP-provided movement regressors from both the LR and RL runs. We then calculated the average FD across all participants, which was low (group mean FD = 0.16 mm). Given the overall low motion in the sample and the extensive motion handling already implemented in the HCP pipeline, no subjects were excluded based on individual FD values.

### 2.4 Striatal Parcellation

We recently developed a method for identifying striosome-like and matrix-like voxels in the human striatum based on their distinct in vivo connectivity profiles (Waugh, Hassan et al. 2022). We use the terms “striosome-like” and “matrix-like” to emphasize that these parcellations are inferential and probabilistic, in contrast with the gold standard for identifying striosome and matrix in tissue, immunohistochemical staining. To parcellate the striatum, we generated composite target masks representing regions with connectivity biases toward one compartment, as established by prior tract-tracing studies in animals and by diffusion tractography in humans (summarized in Waugh, Hassan et al., 2022). Striosome-favoring regions included the posterior orbitofrontal cortex, anterior insula, basolateral amygdala, basal operculum, and posterior temporal fusiform cortex, whereas matrix-favoring regions included the inferior frontal gyrus pars opercularis, primary motor cortex, supplementary motor area, primary somatosensory cortex, and superior parietal cortex.

Striatal parcellation was performed using the FSL tool **probtrackx2**, which allowed us to evaluate the relative connectivity of each striatal voxel to striosome-favoring versus matrix-favoring target masks. Tractography was conducted in each subject’s native diffusion space using standard parameters: curvature threshold = 0.2, step length = 0.5 mm, number of steps per sample = 2,000, number of samples per seed voxel = 5,000, and distance correction enabled to prevent target proximity from biasing connection strength. For each voxel, we compared the number of streamlines projecting to striosome-favoring versus matrix-favoring targets, and the resulting ratio of streamline counts was used as an index of compartmental bias. This produced a probability value for each voxel (P = 0–1), indicating the degree of striosome- or matrix-like connectivity. Voxels were classified as compartment-biased when P > 0.55 toward either set of targets, yielding continuous subject- and hemisphere-specific maps of striosome-like and matrix-like connectivity. Because the precise spatial distribution of striosome is unique to each individual, these parcellations must be defined for each individual; standardized striatal region-of-interest masks are not suitable for studying compartmental organization.

Because the resolution of our diffusion voxels matched the upper limit of the diameter of striosome branches (1.25 mm), all voxels had the potential to contain both striosome and matrix tissue. In fact, many striatal voxels exhibited no or only modest compartment-like connectivity bias. To maximize the contrast between compartments, we excluded voxels with low or indeterminate bias using an iterative thresholding procedure applied to the compartmental probability maps. For each subject and hemisphere, we selected voxels from the most strongly biased portion of the distribution (favoring either striosome or matrix connectivity) and iteratively lowered the threshold until the mask volume reached a predetermined target. This target was set at 13% of the original striatal mask, corresponding to 1.5 standard deviations above the mean of a Gaussian distribution. To ensure equality of mask volumes, which reduced size-based biases, we matched striosome-like and matrix-like mask volume within each subject and hemisphere. We have previously demonstrated that these high-bias, equal-volume masks reproduce the characteristic spatial distribution, differential connectivity, and somatotopy of striosome and matrix observed in histological studies (Waugh, Hassan et al. 2022, Funk, Hassan et al. 2023, Funk, Hassan and Waugh 2024, Sadiq, Funk and Waugh 2025, Sadiq and Waugh 2025, Waugh and Tieu 2025). Because the diffusion data were acquired at 1.25-mm isotropic resolution (voxel volume = 1.953 mm^3^) and the fMRI data at 2-mm isotropic resolution (voxel volume = 8 mm^3^), a given anatomical volume is represented by fewer voxels in fMRI space than in diffusion space. Based on the volumetric ratio of approximately 4.1 (8 ÷ 1.953), 311 diffusion-space voxels translated to roughly 76 voxels at fMRI resolution. This volume-based conversion provides a consistent and biologically grounded reference for comparing compartment sizes across modalities and ensures appropriate interpretation when projecting diffusion-derived masks into fMRI space. Each subject’s high-bias compartment masks were then registered from diffusion space to structural (T1-weighted) space using FSL’s FLIRT, and subsequently nonlinearly transformed into fMRI space with the precomputed functional-to-structural registration matrices from the HCP pipeline. All transformations were visually verified for accuracy. The resulting striosome-like and matrix-like masks in fMRI space were used as seeds in subsequent functional connectivity analyses.

### 2.5 Validation of Striatal Parcellation

To test whether our parcellated voxels reproduced the anatomic patterns of striatal compartmentalization established through histology, we quantified their spatial organization within the equal-volume 1.5 SD masks. For each subject and hemisphere, Cartesian coordinates (x, y, z) were extracted for every voxel and referenced to the centroid of the corresponding nucleus (caudate or putamen). We then assessed spatial distribution by calculating within-plane dispersion and the root-mean-square (RMS) distance from the nucleus centroid, providing voxelwise measures of compartment organization in three-dimensional striatal space. Consistent with prior histological descriptions, we have previously shown that striosome-like voxels are preferentially localized to rostral, medial, and ventral striatum (Graybiel and Ragsdale Jr 1978, Goldman‐Rakic 1982, Donoghue and Herkenham 1986, Ragsdale Jr and Graybiel 1990, Desban, Kemel et al. 1993, Eblen and Graybiel 1995, Waugh, Hassan et al. 2022).

Striosomal branches are embedded within the surrounding matrix (Graybiel and Ragsdale Jr 1978, Holt, Graybiel and Saper 1997). In coronal histologic sections, striosome appear as discrete “islands” within a continuous “sea” of matrix tissue, though the striosome is actually a contiguous, 3-dimensional structure. To compare the spatial organization of our MRI-derived parcellations with this observation from tissue, we applied the *fsl-cluster* command at a high bias threshold (P > 0.87) to isolate voxels with strong compartment-specific bias. We analyzed the largest cluster within each compartment, comparing the degree of separation (contiguity vs. isolation) in striosome- and matrix-like voxels.

We previously showed, in a similar HCP-derived cohort, that shifting voxel location by only 2-3 mm was sufficient to eliminate striosome-like bias in structural connectivity, indicating that compartment-specific biases depend on precise voxel position rather than the striatal “neighborhood” in which a voxel is located (Funk, Hassan et al. 2023). More recently, we demonstrated that shifting voxel location also eliminated compartment-specific biases in resting-state functional connectivity (Sadiq, Funk and Waugh 2025). Based on these findings, we hypothesized that task-evoked functional biases would similarly depend on precise voxel location. To test this, we assessed the spatial specificity of task-based activation by jittering the locations of striosome- and matrix-like voxels by ±0–3 voxels in each anatomical plane, randomly and independently for each voxel. For every subject, we confirmed that although individual voxels were displaced, the mean location of the jittered masks remained nearly identical to the mean of the original masks, and importantly, no jittered voxel overlapped with voxels from the original masks. The average root-mean-square displacement was small: 2.9 voxels for the randomized striosome-like mask and 3.1 voxels for the randomized matrix-like mask. When shifts were averaged within each anatomical plane (combining positive and negative displacements), the mean displacement was minimal (0.37 voxels; range: 0.07– 1.2). Thus, the jittering procedure produced small but measurable displacements at the voxel level without altering the overall spatial neighborhood of the striatal masks. In our prior motor task fMRI study (Sadiq and Waugh 2025), we showed that such jittering was sufficient to abolish compartment-specific activation differences specifically in the striosome compartment, confirming that observed biases were spatially precise and not attributable to local neighborhood effects. Consistent with this framework, we used the jittered masks here as a negative control, directly comparing task-evoked activation in the original striosome- and matrix-like voxels with activation in the randomized voxel location masks.

### 2.6 Analysis of task-based functional MRI (tfMRI)

We performed a task-based fMRI analysis to characterize condition-specific activation within striosome-like and matrix-like striatal compartments during working memory tasks. At the first level, statistical modeling was carried out using a general linear model (GLM), with each working memory condition (0-back and 2-back) represented as a 25-second block defined by onset timings from the HCP-provided explanatory variable (EV) files. To account for acquisition variability, data from left-to-right (LR) and right-to-left (RL) phase-encoding runs were combined in a second-level analysis, yielding averaged estimates. For each subject, contrasts of parameter estimate (COPEs) were extracted for the motor conditions, and mean activation values were calculated within the striosome-like and matrix-like masks. This activation measure corresponded to COPE estimates from the subject-level GLMs and are expressed in arbitrary units (AU), reflecting the relative amplitude of BOLD signal change.

Unlike resting-state functional connectivity, which measures spontaneous correlations among networks, this task-based approach isolates activity that is aligned with specific task events and characterizes how that activity unfolds across time. In doing so, it provides a direct test of how striosome- and matrix-like voxels are differentially recruited across distinct phases of memory performance.

### 2.7 Slice-wise Voxel Sampling for Sensitivity Analysis

Our primary voxel-selection strategy emphasized voxels with the strongest compartmental bias but did not explicitly control for their spatial distribution within the striatum. As a consequence, highly biased voxels, particularly within the matrix-like distribution, could be spatially clustered, raising the possibility that observed compartment-specific effects might reflect regionally concentrated signals rather than striatum-wide organizational principles. Our previously described striosome-like masks utilized all, or nearly all voxels in the striosome-like distribution (target: 13% of striatal volume; expected from tissue: 15% of striatal volume). Re-selecting striosome-like voxels was unlikely to substantially change these masks. In contrast, the matrix-like distribution included many voxels that were unsampled in our primary matrix-like masks, and these tended to be substantially more clustered than striosome-like voxels. To evaluate whether matrix-like effects were robust to this voxel-selection assumption, we performed a sensitivity analysis using a slice-wise voxel sampling approach to define our matrix-like masks.

Specifically, for each axial slice of the striatum, we selected an equal number of the most-biased matrix-like voxels. This procedure ensured that matrix-like voxels were sampled across the full superior–inferior extent of the striatum while preserving our goal of using high-bias voxels to represent each compartment. This approach reduced the influence of localized clustering and tested the sensitivity of compartment-specific activation effects to voxel selection strategy. Persistence of matrix-like effects under this alternative sampling scheme would indicate that these effects reflect striatum-wide compartmental organization rather than localized regional bias.

### 2.8 Statistical Analysis

We assessed the accuracy of our striatal parcellations – the intra-striate position, clustering, and mean bias of our compartment-like voxels – using a series of two-tailed, paired-samples t-tests. We compared the volume within streamline bundles seeded by compartment-like voxels using two-tailed, paired-samples t-tests.

For the task-based analyses, separate comparisons tested (1) compartment-specific activation during cue presentation, (2) load-dependent activation (2-back vs. 0-back), (3) accuracy-related modulation (correct vs. error trials), and (4) category selectivity (each stimulus type vs. the mean of all other stimulus types). To control for multiple comparisons, we applied the Benjamini-Hochberg (BH) procedure (Benjamini and Hochberg 1995) across each family of related tests (e.g., stimulus categories × task loads × compartments), using a false discovery rate (FDR) threshold of Q = 0.05. For transparency, corrected significance thresholds are reported in the Results. We defined a result as trending toward significance if p < 0.05 but the result did not survive multiple comparisons correction.

Because there were very few errors in 0-back trials, we restricted our accuracy-based analyses to the 2-back condition (n = 844 participants with sufficient error trials). All activation values were averaged across hemispheres to yield mean compartment-level activation per subject.

## 3 Results

After applying exclusion criteria, the final sample included 965 healthy adults (mean age = 29.3 ± 3.7 years; 527 females, 438 males). Participants were excluded primarily due to incomplete diffusion or task-fMRI data or due to self-reported substance or alcohol use. This cohort partially overlapped with our previously reported resting-state sample (Sadiq, Funk and Waugh 2025), with 674 participants (70%) common to both analyses.

### 3.1 Comparing MRI-parcellated voxels to striosome and matrix in tissue

To validate our MRI-based parcellations, we first tested whether striosome-like and matrix-like voxels reproduced the well-established spatial organization of these compartments within the striatum. Across species ranging from rodents to primates, striosome branches are consistently enriched in the rostral, ventral, and medial striatum, whereas the matrix compartment predominates in caudal, dorsal, and lateral regions (Graybiel and Ragsdale Jr 1978, Goldman‐Rakic 1982, Donoghue and Herkenham 1986, Ragsdale Jr and Graybiel 1990, Desban, Kemel et al. 1993, Eblen and Graybiel 1995, Waugh, Hassan et al. 2022). Our voxelwise location analyses replicated this bias bilaterally. In the caudate, matrix-like voxels were significantly more lateral (1.3 mm; p = 8.6×10−^52^), caudal (−9.4 mm; p < 1×10^−260^), and dorsal (8.7 mm; p < 1×10^−260^) than striosome-like voxels. Similarly, in the putamen, matrix-like voxels were more lateral (0.7 mm; p = 8.8×10^−5^), caudal (−5.8 mm; p < 1×10^−260^), and dorsal (5.9 mm; p < 1×10^−260^) than striosome-like voxels. Distance-to-centroid RMS analyses confirmed this spatial dissociation: in the caudate, striosome-like voxels clustered 12.2 mm medio-rostro-ventral to the centroid, whereas matrix-like voxels were positioned 13.7 mm latero-caudo-dorsal; in the putamen, striosome-like voxels were 10.2 mm medio-rostro-ventral, compared to 8.0 mm latero-caudo-dorsal for matrix-like voxels.

Histological studies in both animal and human tissue found that striosome and matrix comprise roughly 15% and 85% of the striatal volume, respectively (Johnston, Gerfen et al. 1990, Desban, Kemel et al. 1993, Holt, Graybiel and Saper 1997). To evaluate whether our MRI-based parcellations approximated this ratio, we quantified compartment-like volume across a range of bias thresholds. Because diffusion voxels (1.25 mm isotropic) sample tissue without regard to compartment boundaries, many voxels inevitably contain mixtures of striosome and matrix. Increasing the bias threshold excludes more of these blended voxels, yielding volume estimates based on voxels with stronger compartment-specific connectivity. At the highest cutoff (P > 0.87), striosome-like voxels represented 5.9% of biased voxels and matrix-like voxels 94.1%, while the majority of striatal voxels (82%) remained unclassified (i.e., indeterminate bias). At the lowest cutoff (P > 0.55), 25.5% of biased voxels were striosome-like and 74.5% were matrix-like, with 45% of the striatum showing sufficient compartment-like bias to be classified. The threshold that most closely matched histological estimates (15:85) was P > 0.7, which yielded 17.3% striosome-like voxels and 82.7% matrix-like voxels, encompassing 32% of total striatal volume. Although proportions shifted with the chosen threshold, all analyses demonstrated consistent volumetric relationships (matrix-like much larger than striosome-like) that paralleled histological observations. This correspondence supports the validity of our MRI-based parcellation approach for identifying compartment-like organization in vivo.

Finally, we examined whether striosome-like and matrix-like voxels differed in their tendency to cluster with other voxels of the same type. In histologic sections, each striosome branch is surrounded by contiguous matrix tissue (Graybiel and Ragsdale Jr 1978, Holt, Graybiel and Saper 1997). In contrast, diffusion MRI samples the striatum using a rigid voxel grid that does not conform to individual striosome architecture, introducing partial volume effects that blur striosome-like signal across adjacent voxels. Although this resolution is insufficient to resolve the connectivity of individual striosome branches, we evaluated whether striosome- and matrix-like voxels differed in their clustering tendencies. Striosome-like voxels tended to form smaller, more scattered clusters, whereas matrix-like voxels aggregated into larger, more contiguous clusters. In the right hemisphere, matrix-like clusters were on average 4.2 times larger than striosome-like clusters, and in the left hemisphere they were 3.4 times larger. In the right hemisphere, striosome-like clusters had a mean volume of 234.2 mm^3^ [SEM ± 6.1; 95% CI: 222–246], substantially smaller than the mean matrix-like cluster volume of 985 mm^3^ [SEM ± 10.9; 95% CI: 963–1006]; p = 1.5×10^−275^. Similarly, in the left hemisphere, striosome-like clusters averaged 301 mm^3^ [SEM ± 6.7; 95% CI: 287–314], compared to 1009 mm^3^ [SEM ± 12.4; 95% CI: 984–1033] for matrix-like clusters; p = 6.0×10^−226^. These results parallel histological observations, in which striosome branches appear as spatially dispersed islands embedded within the continuous matrix.

### 3.2 Cue-Evoked Compartment-Specific Activation in 0-Back and 2-Back Tasks

We assessed cue-specific activation in striosome- and matrix-like voxels during the 0-back and 2-back tasks (Figure 2). During cue periods in the 0-back condition, striosome-like voxels exhibited significantly greater activation than matrix-like voxels across all stimulus categories (body part, face, place, and tool), indicating a consistent compartment-like bias during task preparation. In the 2-back condition, cue-related striosome-like activation remained higher than matrix-like activation for body part and face cues, though compartment differences were no longer significant for place and tool stimuli. Overall, cue-evoked activation across all categories was markedly higher in striosome-like voxels than in matrix-like voxels (Figure 2A). When averaged across categories, striosome-like activation was 1.5-fold greater than matrix-like activation (striosome-like = 40.8, matrix-like = 26.4; p = 1.5×10–^6^). In the 2-back condition, a compartment effects during the cue period was observed only for body part and face stimuli, where striosome-like activation exceeded matrix-like activation ([striosomebody = 62.5, matrixbody = 27.8; p = 6.5*x*10–^8^], [striosomeface = 64.3, matrixface = 51.5; p = 2.3×10^−2^], Figure 2B). Multiple comparisons correction yielded a corrected significance threshold of p = 3.8×10^−2^.

**FIGURE 2:**
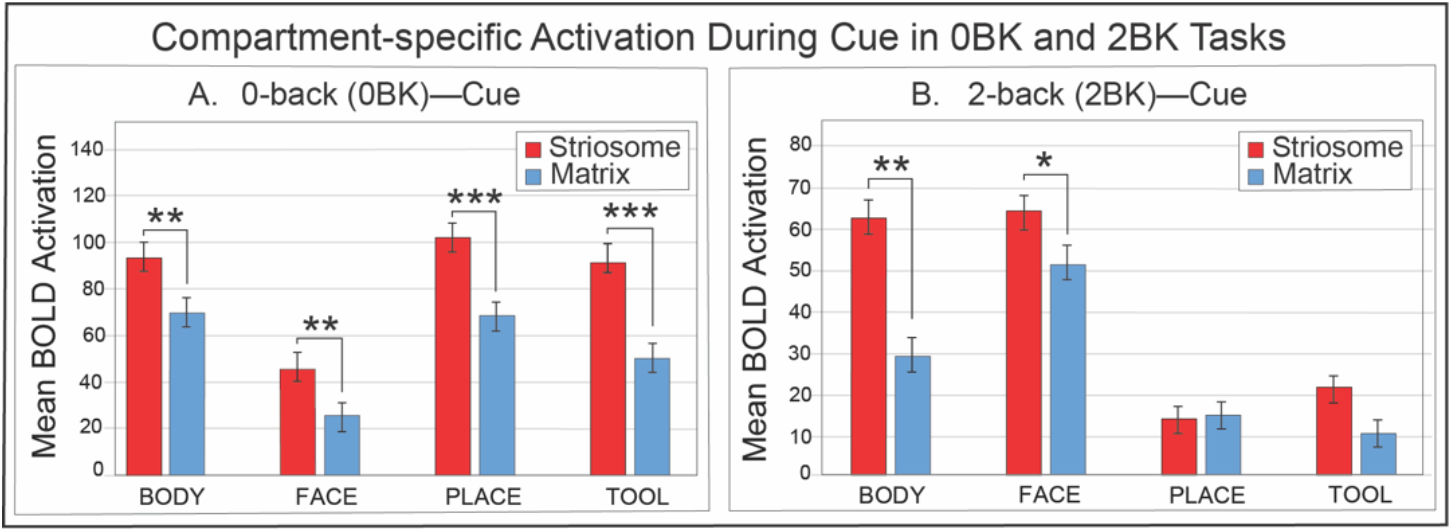
Compartment-specific activation during the cue phase of the 0-back and 2-back working memory tasks. Mean BOLD activation is plotted for striosome-like (red) and matrix-like (blue) voxels across four stimulus categories (body part, face, place, tool) during the cue period. (A) In the 0-back task, striosome-like voxels showed significantly greater activation than matrix-like voxels during cues for all categories, indicating strong cue-related engagement of the striosome across stimulus domains. (B) In the 2-back task, striosome-like activation remained higher than matrix-like activation during body part and face cues, whereas no significant differences were observed for place and tool cues. These findings suggest that striosome-like voxels are preferentially recruited during anticipatory stages of working memory tasks, particularly for socially and biologically salient categories, while matrix-like voxels exhibit weaker cue-related responses. Error bars illustrate the standard error of the mean. *, p < 0.05; **, p < 1×10^−8^; ***, p < 1×10^−^10.

### 3.3 Compartment-Specific Activation Patterns During Working-Memory Tasks

Next, we assessed the post-cue activation in striosome- and matrix-like voxels (Figure 3). Matrix-like voxels exhibited significantly greater activation during working memory tasks, with differences becoming more pronounced under higher cognitive load (2-back). In the 0-back condition, a compartment effect was observed only for *face* stimuli, where matrix-like activation exceeded striosome-like activation (matrix = 11.6, striosome-like = 6.6; p = 4.1×10??; Fig. 3A). In the 2-back condition, however, activation across all stimulus categories was consistently higher in matrix-like voxels (Fig. 3B). When averaged across categories for 2BK, matrix-like activation was 1.3-fold greater than striosome-like activation (matrix-like = 16.6, striosome-like = 12.4; p = 1.9×10^−19^). Multiple comparisons correction yielded a corrected significance threshold of p = 3.1×10^−2^.

**FIGURE 3:**
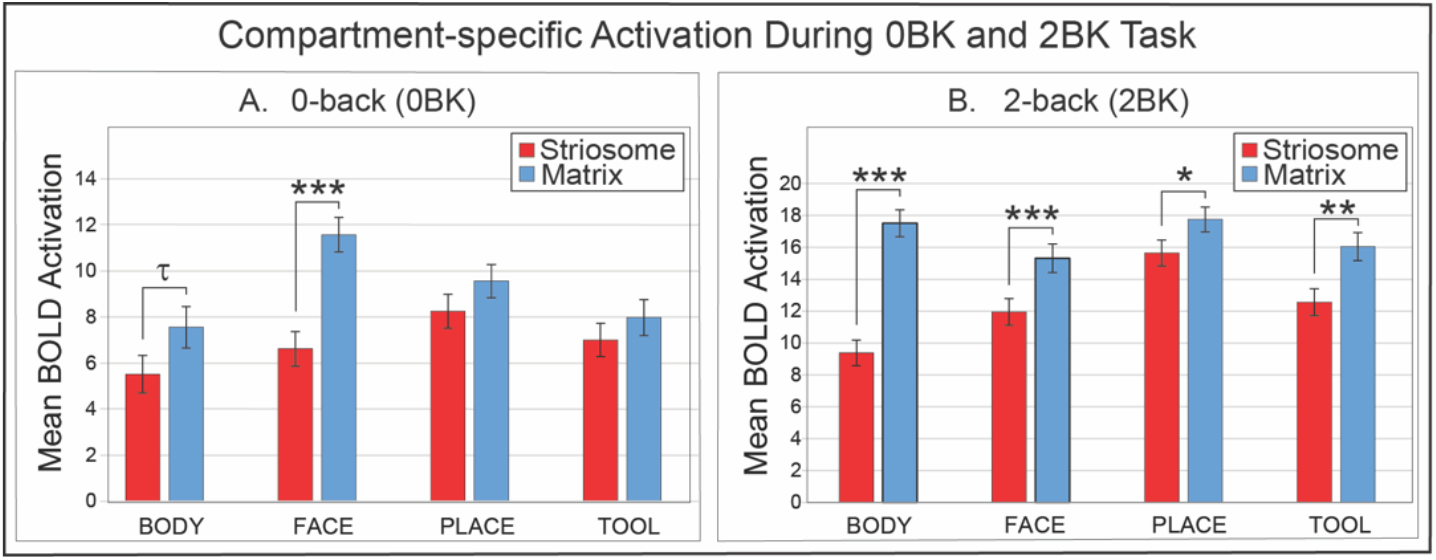
Compartment-specific activation during 0-back and 2-back working memory task blocks. Mean BOLD activation is shown for striosome-like (red) and matrix-like (blue) voxels across four stimulus categories (body part, face, place, tool). (A) In the 0-back condition, matrix-like voxels exhibited significantly greater activation than striosome-like voxels for face stimuli, while no differences were observed for the other categories. (B) In the 2-back condition, matrix-like voxels showed consistently stronger activation than striosome-like voxels across all stimulus categories, with significant effects for body part, face, place, and tool stimuli. This load-dependent divergence suggests that matrix-like voxels are preferentially engaged during the sustained demands of working memory, while striosome-like voxels remain less responsive during active task execution. Error bars illustrate the standard error of the mean. *, p < 0.031; **, p < 1×10^−5^; ***, p < 1×10^−10^; τ, trend toward significance.

While the direction of compartment-specific effects was consistent across load conditions, activation magnitude showed opposite scaling across task epochs. Execution-related activation increased from the 0-back to the 2-back condition, whereas cue-locked activation was markedly larger in the 0-back condition and reduced under higher load.

### 3.4 Sensitivity Analysis Using Less-Biased Voxels

Before evaluating task-evoked effects, we first characterized how the slice-wise voxel sampling procedure altered the properties of the matrix-like masks relative to the primary equal volume 1.5 SD masks. As intended, the slice-wise approach produced matrix-like masks with a modest but consistent reduction in compartmental bias, with mean bias decreasing from 0.955 in the 1.5 SD masks to 0.950 in the slice-wise masks (0.48% decrease). This confirms that the slice-wise masks were less dominated by the most extreme matrix-biased voxels while still retaining strong compartmental specificity.

More importantly, the slice-wise procedure substantially altered the spatial organization of matrix-like voxels. Average cluster volume was reduced from 143.9 to 46.3 voxels, a 67.8% decrease, and the average number of clusters decreased from 14.5 to 6.5 (54.7% decrease). These changes indicate a marked reduction in localized voxel clustering and a redistribution of matrix-like voxels across the striatum, demonstrating that the slice-wise masks differed meaningfully from the primary masks in both bias strength and spatial topology.

Despite these substantial changes in voxel selection and spatial organization, the compartmental dissociations observed in the primary analysis were largely preserved. Matrix-like voxels continued to show greater activation during task execution, whereas striosome-like voxels exhibited stronger cue-period activation. These effects were most robust for body- and face-related conditions across both 0-back and 2-back tasks.

Several contrasts, most notably those involving place stimuli did not retain statistical significance under the slice-wise voxel definition, and in a small subset of conditions (e.g., 2-back place execution and cue), effect directions were not preserved. This pattern suggests reduced robustness rather than systematic reversal and indicates that place-related compartmental effects are more sensitive to the intra-striate location of matrix-like voxels.

### 3.5 Impact of specific voxel location on striatal compartmentalization

Striosome and matrix compartments are distributed in distinct regions of the striatum, and our striosome-like and matrix-like voxel masks showed a similar spatial distribution. This raises the possibility that the observed compartment-specific differences in functional activation might not reflect intrinsic properties of the compartments themselves, but instead the activation patterns of the striatal territories where those voxels were predominantly located—a potential “neighborhood effect.” To address this concern, we compared activation values obtained from the original striosome- and matrix-like masks (Section 3.2) with those derived from spatially shifted versions of the same masks, thereby testing whether compartment-specific effects depended on the precise voxel locations selected. We applied voxelwise jittering to each compartment-like mask, such that the position of individual voxels was randomized while the overall displacement of the mask remained minimal. On average, the center of gravity of the jittered masks shifted only 0.37 voxels from the original masks. In matrix-like voxels, mean bias decreased from 0.95 in the original mask to 0.83 in the jittered version (p < 1×10^−260^). In striosome-like voxels, bias declined more sharply, from 0.78 to neutral (0.47; p < 1×10^−260^).

These findings indicate that compartment-specific bias cannot be attributed to regional location (neighborhood effects) but instead reflected the structural connectivity of the precisely defined voxels within our compartment-like masks. We therefore set these location-shifted voxels as negative controls to assess the specificity of compartment-specific activation during working memory. In spatially shifted striosome-like masks, functional activation during both 0-back and 2-back working memory tasks was reduced across all eight task conditions, compared with the original, precisely-selected striosome-like masks (Table 1). In the 0-back condition, shifting voxel locations reduced activation for all stimulus categories, with decreases ranging from −8.9% (*place*, p = 2.5×10^−2^) to −32.1% (*body* part, p = 2.5×10^−6^). In the 2-back condition, activation was again reduced across all categories, with reductions of −13.6% (*place*, p = 2.0×10^−9^) to −30.8% (*body* part, p = 5.0×10^−14^). These findings demonstrate that even when location-shifted voxels occupied the same striatal “neighborhood,” their striosome-like activation profile was critically dependent on precise voxel placement.

**Table 1:** Randomly shifting the location of striosome-like voxels by 2–3 voxels significantly reduced functional activation during working-memory tasks, whereas shifting matrix-like voxels had minimal effects. This matches the expectation from histology: since striosome is surrounded by matrix, shifting from striosome-like voxels is likely to select a matrix-like voxel and thus change the pattern of activation. In contrast, shifting the location of a matrix-like voxel is likely to select another matrix-like voxel. Δ Activation (Shifted – Original), percentage change, and p-values indicate how voxel displacement altered mean activation within each compartment. Bolded p-values denote significant differences (corrected for multiple comparisons).

| Compartment | Task | $\Delta$ Activation<br>(Shifted – Original) | % Change | *p*-value |
| --- | --- | --- | --- | --- |
| 0-back<br>Striosome | *Body part* | -1.8 | -32.1 | **$2.5 \times 10^{-6}$** |
|  | *Face* | -2.1 | -31.6 | **$4.9 \times 10^{-9}$** |
|  | *Place* | -0.7 | -8.9 | **$2.5 \times 10^{-2}$** |
|  | *Tool* | -1.8 | -25.9 | **$9.9 \times 10^{-8}$** |
| 0-back<br>Matrix | *Body part* | -0.4 | -5.3 | $2.7 \times 10^{-1}$ |
| | *Face* | -0.3 | -2.3 | $4.5 \times 10^{-1}$ |
| | *Place* | -0.4 | -4.6 | $1.7 \times 10^{-1}$ |
| | *Tool* | -0.2 | -2.0 | $6.2 \times 10^{-1}$ |
| 2-back<br>Striosome | *Body part* | -2.8 | -30.8 | **$5.0 \times 10^{-14}$** |
|  | *Face* | -2.7 | -22.3 | **$1.1 \times 10^{-13}$** |
|  | *Place* | -2.1 | -13.6 | **$2.0 \times 10^{-9}$** |
|  | *Tool* | -3.5 | -27.7 | **$1.1 \times 10^{-18}$** |
| 2-back<br>Matrix | *Body part* | -0.3 | -1.9 | $3.7 \times 10^{-1}$ |
| | *Face* | -0.73 | -4.8 | $3.8 \times 10^{-2}$ |
| | *Place* | -0.2 | -1.4 | $5.0 \times 10^{-1}$ |
|  | *Tool* | -1.2 | -8 | **$9.7 \times 10^{-4}$** |

In the matrix-like compartment, shifting voxel locations had minimal impact on activation. For the 0-back condition, activation changes were uniformly small (−2.0% to −5.3%) and non-significant across all stimulus categories (all p ≥ 0.17). For the 2-back condition, reductions were similarly minimal (−1.4% to −4.8%), except for the *tool* category, which showed a significant decrease (−8.0%, p = 9.7×10^−4^). This stability likely reflects the abundance and widespread distribution of matrix-like voxels, such that shifting the location of a matrix-like voxel most often results in selecting another matrix-like voxel with slightly weaker bias. This control analysis highlights the biological specificity of compartment-related differences: shifting striosome-like voxels removes their striosome-like properties, whereas shifting matrix-like voxels only reduces the strength of their matrix bias. Thus, compartment-specific effects reflect precise voxel selection rather than nonspecific activation patterns of the surrounding striatal “neighborhood. Notably, following voxel displacement all activation differences were negative, indicating that voxels with stronger structural connectivity bias also exhibited stronger functional activation, further supporting a tight coupling between structural compartmentalization and functional engagement.

### 3.6 Activation Changes from 0-Back to 2-Back Within Striatal Compartments

Next, we investigated within-compartment differences between the 0-back and 2-back conditions to assess how increasing working memory load modulated striatal activation. During task execution, both striosome- and matrix-like voxels exhibited significantly higher activation during the 2-back condition compared to the 0-back condition (Figure 4). Both striosome- and matrix-like voxels showed robust increases in activation with the increased effort of the 2-back condition, with fold-changes ranging from ∼1.7-to 1.9-fold in the striosome-like and ∼1.3-to 2.3-fold in the matrix-like voxels, reflecting proportional increases in mean BOLD signal within each compartment (Figure 2). Specifically, striosome-like activation rose by 1.7× for body part, 1.8× for face, 1.9× for place, and 1.8× for tool stimuli, while matrix-like activation increased by 2.3×, 1.3×, 1.8×, and 2.0× for the same categories, respectively. Since all comparisons survived correction, the effective corrected significance threshold was p < 0.05.

**FIGURE 4:**
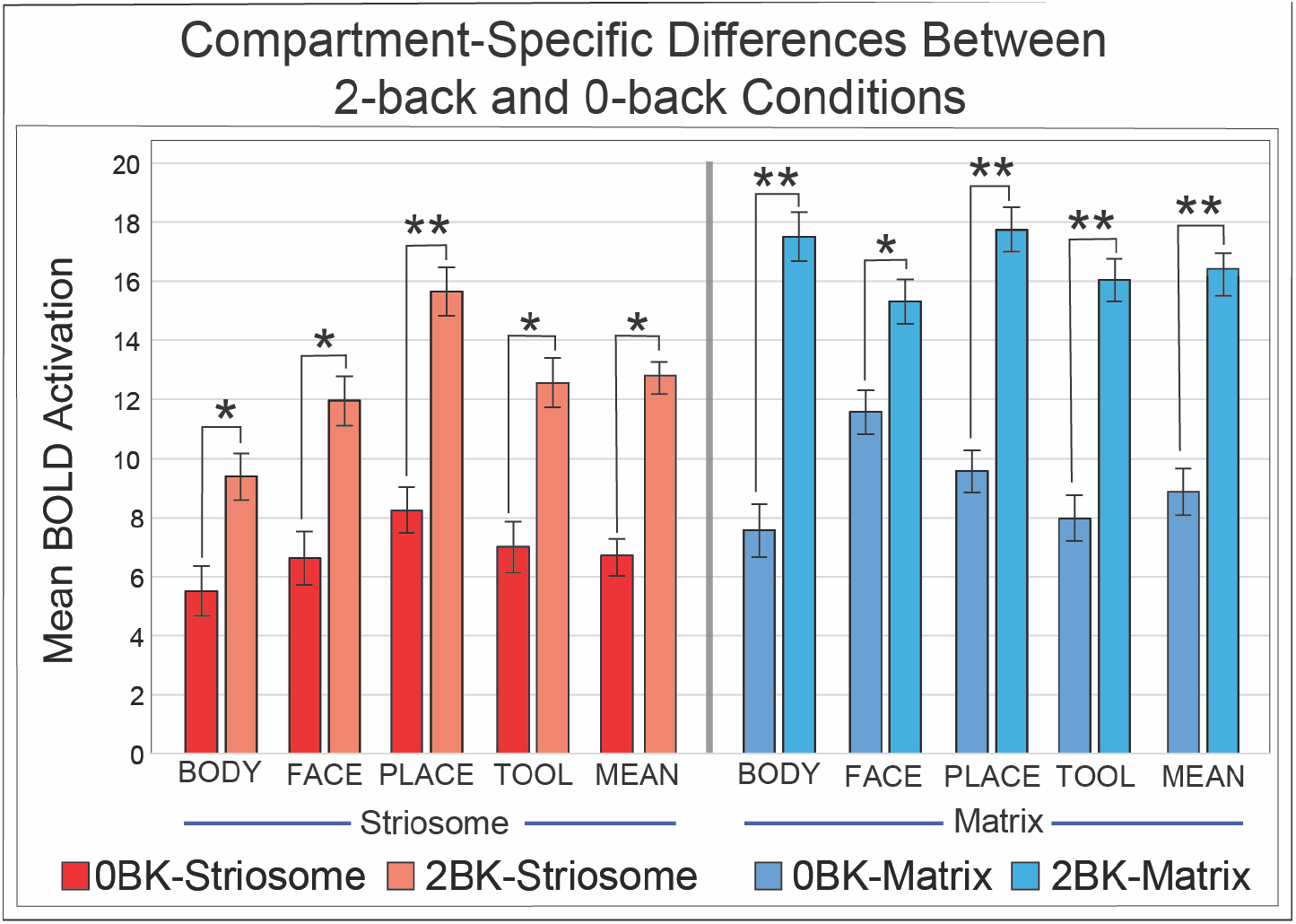
Activation in 2-back memory tasks was higher for every condition, in both compartments, than activation in 0-back tasks. Mean BOLD activation is shown separately for striosome-like (red/orange) and matrix-like (blue shades) voxels across four stimulus categories (*body* part, *face, place, tool*). For both compartments, activation increased significantly in the 2-back relative to the 0-back condition, indicating sensitivity to working memory load. Striosome-like voxels (left) showed modest but reliable increases for *body* part, *face, place*, and *tool* stimuli, suggesting a limited but consistent contribution under higher task demands. In contrast, matrix-like voxels (right) exhibited much stronger load-dependent increases across all categories, with the largest effects observed for *body* part and *place* stimuli. These findings demonstrate that while both compartments respond to increasing cognitive load, the matrix shows greater scaling of activation with task difficulty, consistent with its role in sustaining execution processes. Error bars illustrate standard error of the mean. *, p < 0.05; **, p < 1×10^−8^.

### 3.7 Compartment-specific Activation in Correct and Error Trials

We next investigated whether trial accuracy correlated with compartment-specific task activation by comparing correct and error trials (Figure 5). Notably, we assessed activation differently in this experiment – unlike other analyses that showed overall or stimulus-locked task activation, this analysis estimated activity that was uniquely attributable to specific trial types relative to the implicit baseline. Analyses were limited to participants with sufficient data for each condition: all subjects contributed 0-back correct trials (n = 965), but very few subjects made errors in this simple task. In contrast, most participants had both correct and incorrect responses in the 2-back trials (n = 844). Activation values for correct and error trials were analyzed separately rather than as difference scores (as in our other analyses), allowing direct comparison of mean BOLD responses between accuracy conditions within each compartment. Correct trials were available for both the 0-back and 2-back conditions and were aggregated across all stimulus categories (*body, face, place*, and *tool*). Since 0-back error trials were absent for most participants, we examined error-related activation only for the 2-back condition. Matrix-like voxels exhibited significantly greater activation than striosome-like voxels during correct trials in both the 0-back and 2-back tasks (Fig. 5A). In the 0-back correct condition, matrix-like activation was 17.9-fold higher than striosome-like activation (matrix: 6.3, striosome: 0.35; p = 5.6×10^−33^). Because sustained task-related engagement during the 0-back condition was present throughout the block and was not uniquely time-locked to individual correct events, this ongoing activity was absorbed into the baseline. As a result, striosome responses during 0-back correct trials were near zero. That is, the robust task-related activation observed in striosome-like voxels in our other analyses was sustained in 0-back trials, but did not increase further in correct trials. The difference between compartments was amplified during the 2-back correct condition, where matrix-like activation nearly doubled striosome-like activation (matrix: 14.5, striosome: 8.0; p = 1.3×10^−30^). Importantly, even during error trials in the 2-back task, matrix-like activation remained significantly greater than striosome-like activation (matrix: 10.5, striosome: 2.4; p = 4.5×10^−13^).

**FIGURE 5:**
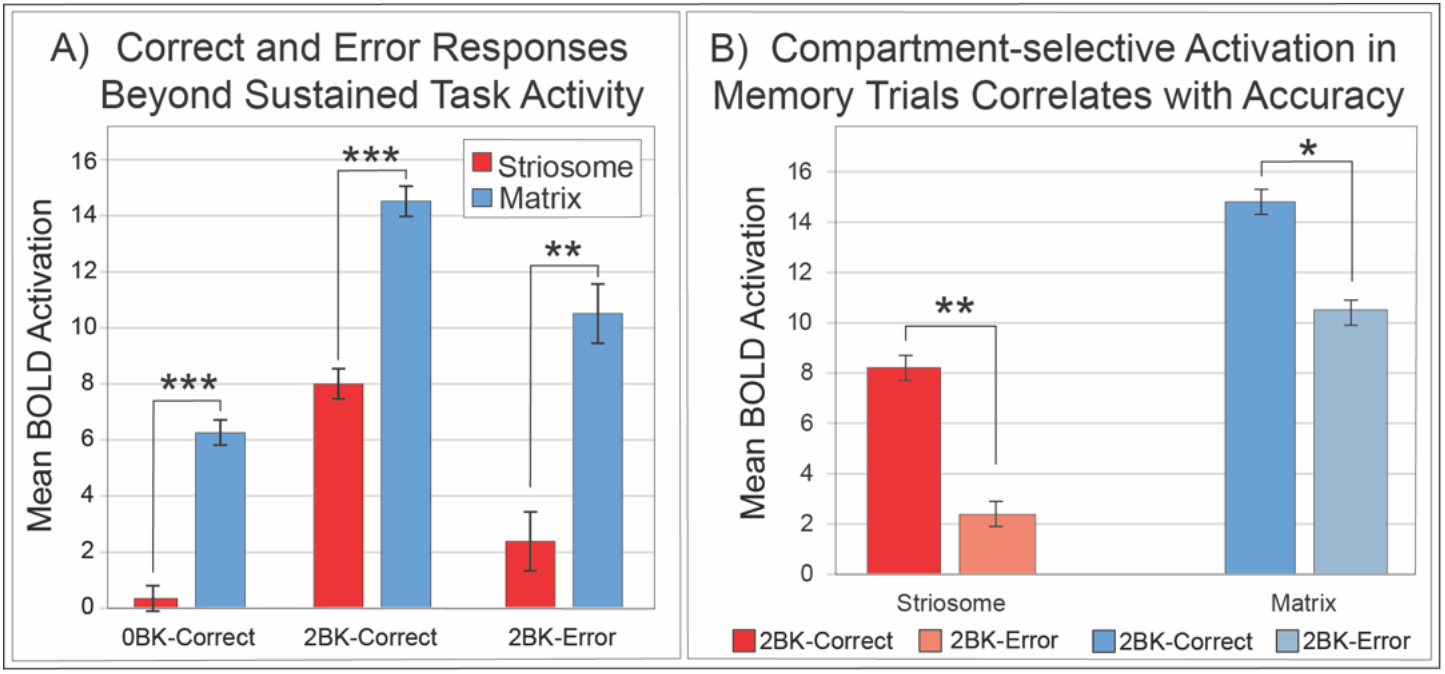
Compartment-specific activation during Correct and Error trials *beyond sustained task activity* in the 0-back and 2-back tasks. (**A)** Mean BOLD activation in striosome-like (red) and matrix-like (blue) voxels during Correct 0-back trials (0BK-Correct), Correct 2-back trials (2BK-Correct), and Error 2-back trials (2BK-Error). Note that most subjects made no errors in the 0-back task, leading to an insufficient sample to assess the 0BK-Error condition. Matrix-like voxels exhibited significantly greater activation than striosome-like voxels in both Correct conditions, with activation levels scaling strongly with task difficulty (2-back > 0-back). Striosome-like voxels also showed increased activation in Correct 2-back trials relative to 0-back, though responses remained consistently lower than those of matrix voxels. During Error trials in the 2-back condition, overall activation was reduced in both compartments but was substantially more reduced in striosome-like voxels. (**B)** The same 2-back data from A, replotted to illustrate the impact of accuracy on within-compartment comparisons of BOLD activation. Both compartments showed reduced activation during Error trials; however, the reduction was substantially larger in striosome-like voxels than in matrix-like voxels. Error bars represent the standard error of the mean. *, p < 1×10^−6^; **, p < 1×10^−*9*^; ***, p < 1×10^−30^.

To determine how trial accuracy influenced activation, we compared correct and error trials within each compartment (Fig. 5B). Analyses were restricted to the 844 participants who had valid activation estimates for both correct and error trials in the 2-back condition. Both compartments showed reduced activation during error trials, but the magnitude of this reduction differed between the compartments. In striosome-like voxels, mean activation declined sharply in error trials, from 8.2 to 2.4 (Correct to Error, 71% reduction; *p* = 3.7×10??), whereas in matrix-like voxels, activation decreased from 14.8 to 10.5 (29% reduction; *p* = 1.1×10^−5^). Although both effects were significant, the larger reduction in striosome-like voxels suggests that accuracy correlated more strongly with striosomal activity than with matrix activity. This pattern suggests that trial success is particularly sensitive to striosomal engagement and that striosomal engagement may contribute to the evaluative or “vigilance” processes required for accurate performance.

### 3.8 Category-specific Activation During 0-back and 2-back Tasks in Striosome-like and Matrix-like Voxels

Next, we evaluated whether category-specific information was preserved within striosome-like and matrix-like voxels as working-memory load increased. This analysis aimed to determine whether compartmental specialization extends beyond temporal differences (cue vs. execution) to include content selectivity, that is, whether each compartment retains distinct patterns of activation for specific visual categories under varying task demands.

For each visual category (*body* part, *face, place, tool*), we compared activation during that category with the mean activation across the other three categories (hereafter, MeanOther; Figure 6). At low load (0-back), striosome-like voxels showed reduced *body* and increased *place* activation, relative to MeanOther ([body: 5.5; MeanOther: 7.3; p < 0.01], [place: 8.3, MeanOther: 6.4; p < 0.01]), with *face* and *tool* not differing significantly. Matrix-like voxels showed reduced *body* and increased *face* activation relative to MeanOther ([body: 7.5, MeanOther: 9.7; p < 0.01], [face: 11.5, MeanOther: 8.4; p < 4.3×10^−5^]), with *place* not significantly different and *tool* trending toward a significant difference. Thus, at 0-back, *body* responses were suppressed in both compartments but the compartments differed in the stimulus categories that were enhanced.

**FIGURE 6:**
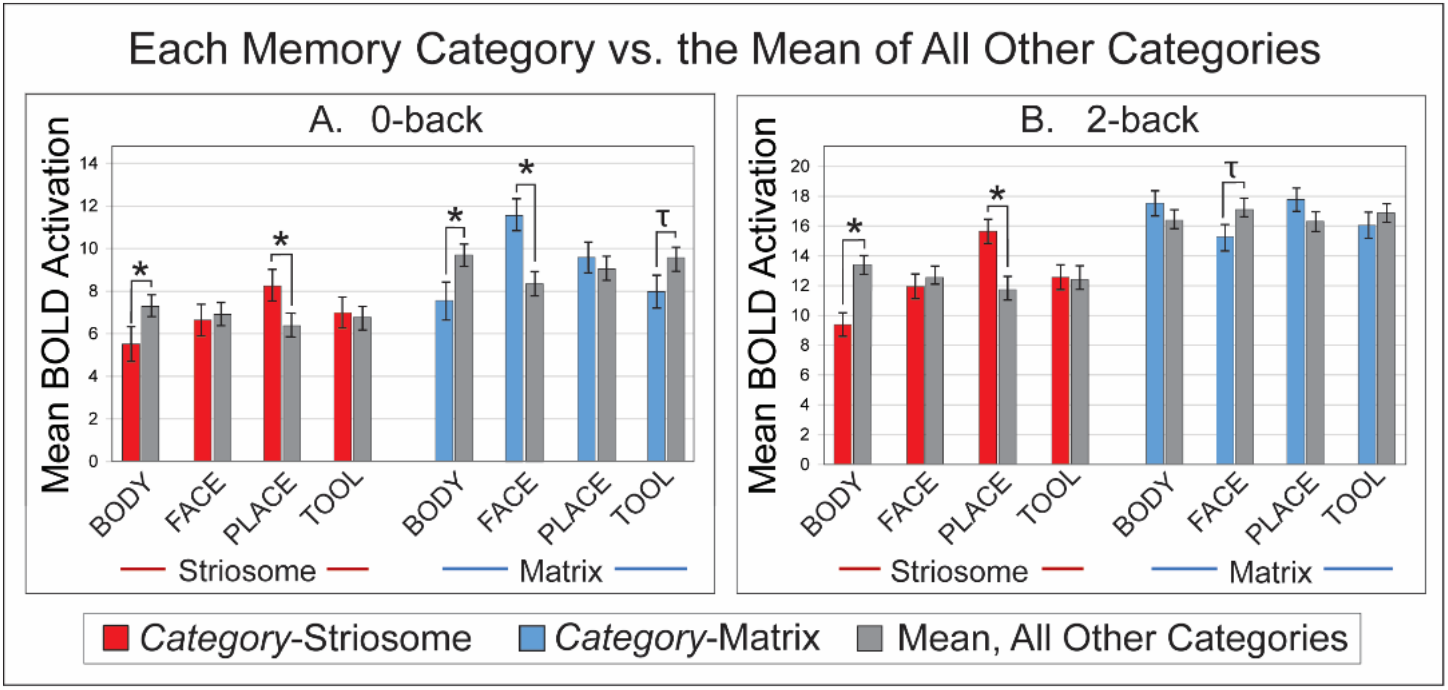
Category-specific activation relative to the mean of all other categories in striosome-like and matrix-like voxels during 0-back and 2-back tasks. Mean BOLD activation is shown for each stimulus category (body part, face, place, tool) relative to the average of the remaining three categories, plotted separately for striosome-like (red) and matrix-like (blue) voxels. (A) **0-back task**. Matrix-like voxels exhibited significant category selectivity, with higher activation for body part, face, and tool stimuli relative to the mean of other categories. Striosome-like voxels showed more limited category selectivity, with significant effects observed for a subset of categories. (B) **2-back task** Striosome-like voxels displayed pronounced category selectivity, with significantly higher activation for body part and place stimuli relative to the other categories, whereas matrix-like voxels showed more uniform responses across categories. Together, these results indicate that both striosome-like and matrix-like voxels exhibit category selectivity under low cognitive demand, whereas increasing task load is associated with a redistribution of category tuning, such that striosome-like voxels remained category-selective but matrix-like voxels lost selectivity. Error bars illustrate the standard error of the mean. *, p < 0.022; τ, trend toward significance.

At higher load (2-back), category modulation became indistinct in matrix-like voxels but enhanced in striosome-like voxels. These findings indicate a compartmental dissociation: under low demand, both compartments show some category sensitivity, but as cognitive load increases, striosome-like voxels maintain category-specific tuning ([body: 9.4; MeanOther: 13.4; p < 3×10^−8^], [place: 15.6, MeanOther: 11.3; p < 2.8×10^−8^]) while matrix-like responses shift to general task engagement. Multiple comparison correction yielded a significance threshold of p = 2.2×10^−2^.

## 4 Discussion

Differences in development, pharmacology, and connectivity predict distinct functions for striosome and matrix (Graybiel and Ragsdale Jr 1978, Crittenden and Graybiel 2011, Brimblecombe and Cragg 2017). Animal studies support compartment-specific roles across valuation/threat decisions, habit learning, and motor control (Friedman, Homma et al. 2015, Xiao, Deng et al. 2020, Nadel, Pawelko et al. 2021, Okunomiya, Watanabe et al. 2025). In humans, evidence for compartment-specific functionality has been largely inferential, based on neuroimaging studies demonstrating distinct striatal connectivity and functional organization consistent with striosome– matrix architecture. For example, a recent study of neuroleptic-induced dystonia reported differential dopaminergic alterations within striosome and matrix territories, suggesting that disruption of their balance contributes to abnormal motor control (Goto 2025). In a recent postmortem investigation of patients with prolonged anorexia nervosa, Kawakami et al., (2022) identified prominent astrogliosis within the striosome of the nucleus accumbens shell, accompanied by neuronal deformation and evidence of impaired dopaminergic innervation between the ventral tegmental area and striatal reward compartments (Kawakami, Iritani et al. 2022). These findings suggest that disruptions in striosome-linked reward processing may contribute to the cognitive rigidity and motivational deficits characteristic of anorexia nervosa. Building on our prior work, which described compartment-specific intrinsic functional networks at rest and a temporal dissociation of compartment activation during motor behaviors (Sadiq, Funk and Waugh 2025, Sadiq and Waugh 2025), we asked whether this striatal compartment specialization extends to cognition.

In the n-back task, striosome-like voxels preferentially activated during preparatory/cue-related demands, whereas matrix-like voxels were more engaged during on-task maintenance and response execution (0-back/2-back blocks). This echoes our motor-task results, where striosome-like voxels were preferentially active during motor preparation and matrix-like voxels during movement execution (Sadiq and Waugh 2025), and complements our resting-state finding that the two compartments occupy distinct intrinsic functional networks (Sadiq, Funk and Waugh 2025). Together, these converging observations indicate that compartment-based specialization is expressed both at rest and dynamically during behavior. In a parallel structural connectivity study, we recently demonstrated that CA1, the primary output node for the hippocampus, has regional biases toward the compartments: rostro-medial CA1 is biased toward matrix-like voxels, while caudo-lateral CA1 is biased toward striosome-like voxels (Tieu, Sadiq et al. 2026). These findings are consistent with animal literature that implicated the striosome in threat valuation, decision making and behavioral inhibition, and implicated the matrix in sensorimotor processing and channeling cortical input into appropriate motor outputs, thereby supporting action initiation (Graybiel 2008, Hikosaka, Kim et al. 2014, Friedman, Homma et al. 2015, Okunomiya, Watanabe et al. 2025). This framework suggests that imbalances in compartment-specific engagement may contribute to motor and cognitive symptoms in neuropsychiatric disease (Tippett, Waldvogel et al. 2007, Crittenden and Graybiel 2011, Waugh, Hassan et al. 2025, Waugh and Tieu 2025).

It is important to acknowledge several limitations of this study. Our method identified striosome- and matrix-like voxels indirectly, based on biases in structural connectivity derived from probabilistic tractography. Consequently, the validity of our functional analyses is contingent upon the anatomical precision of these parcellations. Previous work has shown that our approach to striatal parcellation is highly reliable, with a test-retest error rate of 0.14% (Waugh, Hassan et al. 2022), and that compartment-like biases are highly spatially specific: shifting voxel locations by only a few millimeters eliminates all compartment-like bias in both structural (Funk, Hassan and Waugh 2024, Sadiq, Funk and Waugh 2025, Sadiq and Waugh 2025) connectivity. Furthermore, these connectivity-based parcellations reproduce key anatomical features observed in human and animal tissue, including the relative abundance, spatial distribution, contiguity, and extra-striatal connectivity of striosome and matrix (Waugh, Hassan et al. 2022, Funk, Hassan et al. 2023, Funk, Hassan and Waugh 2024). Nonetheless, this technique cannot substitute for direct histological identification of striosome and matrix. While voxels parcellated through differential connectivity share all of the anatomic properties of striosome and matrix identified in tissue, the extent to which compartment-like voxels match the location of the underlying tissue compartments remains untested.

Another limitation concerns the spatial resolution of diffusion MRI. Although the voxel size used in this study (1.25 mm isotropic) approximates the diameter of the largest human striosome branches (Graybiel and Ragsdale Jr 1978, Holt, Graybiel and Saper 1997), each striosome-like voxel inevitably contains some proportion of matrix tissue. Nevertheless, our previously reported validation studies demonstrated that compartment-like voxels reproduce the anatomical features of striosome and matrix identified through immunohistochemistry, even when derived from diffusion datasets with lower resolution than employed here (Waugh, Hassan et al. 2022, Funk, Hassan et al. 2023, Funk, Hassan and Waugh 2024). The fMRI dataset used here has a comparable resolution to those prior diffusion MRI datasets (2 mm isotropic), and we have shown that this resolution is sufficient to detect robust, widespread compartment-specific patterns of functional connectivity (Sadiq, Funk and Waugh 2025, Sadiq and Waugh 2025). Still, it is important to emphasize that these non-invasive, inferential methods cannot achieve the fine-grained histological detail available in post-mortem tissue. This limitation applies broadly across all current in vivo anatomical mapping approaches in humans.

Since the architecture and precise location of the striosome differs between individuals, standard region-of-interest approaches are inadequate to assess the striatal compartments in vivo. Therefore, we quantified mean activation within striosome-like and matrix-like masks that were uniquely generated for each subject, rather than performing voxel-wise activation analyses. Although averaging within compartments facilitated meaningful group-level comparisons, this method may have obscured more spatially restricted activation patterns. Given that corticostriatal projections are somatotopically organized (Flaherty and Graybiel 1993, Waugh, Hassan et al. 2022, Sadiq, Funk and Waugh 2025, Sadiq and Waugh 2025), it is possible that the functional specializations we observed are confined to specific subregions within each compartment, and do not generalize to the whole striosome, or the whole matrix.

It is also important to acknowledge potential distortions introduced by our experimental design in assessing functional connectivity. Histological studies consistently show that the matrix occupies roughly six times more volume than the striosome (Desban, Kemel et al. 1993, Holt, Graybiel and Saper 1997). To avoid size-related biases in probabilistic connectivity analyses, we generated equal-volume masks, which required assessing only a subset of matrix-like voxels. Consequently, matrix-like voxels outside the most strongly biased subset may exhibit functional activation patterns not captured in our analysis. While this natural volume asymmetry between striosome and matrix provides essential anatomical context, it does not alter the robustness of the functional activation patterns observed in the present study.

These experiments revealed a clear functional dissociation between striosome-like and matrix-like voxels during the cue period. Across both 0-back and 2-back tasks, striosome-like voxels consistently showed stronger activation than matrix-like voxels, suggesting that the striosome may play a specialized role in anticipatory processes related to working memory. Such processes may include attentional orienting, evaluating potential outcomes, and assessing the motivational or emotional significance of upcoming events, all of which align with functions previously attributed to the striosome (Friedman, Homma et al. 2017, Karunakaran, Amemori et al. 2021). Importantly, this compartment bias was evident across multiple stimulus categories (*body* part, *face, place*, and *tool*), highlighting the domain-general nature of striosome-like engagement during task preparation. In contrast, matrix-like voxels showed lower cue-related activity, consistent with their stronger association with execution phases in motor tasks. Together, these findings suggest that during working memory, striosome-like voxels preferentially support cue-dependent processes that establish task context and readiness, while matrix-like voxels may contribute more prominently during subsequent response implementation.

Interestingly, we observed a striking reversal of the compartmental dissociation seen during cue processing. During the active task blocks, matrix-like voxels exhibited significantly greater activation than striosome-like voxels across nearly all stimulus categories, with especially robust differences for *body* part, *face*, and *tool* conditions. This reversal in relative amplitude indicates that the matrix compartment is preferentially engaged once participants transition from preparatory to execution phases, consistent with the proposed role for the matrix in sustaining online processing demands (Flaherty and Graybiel 1993). By contrast, striosome-like voxels showed relatively modest activity during task blocks, supporting the interpretation that their primary contribution lies in anticipatory or evaluative processes rather than in ongoing task execution.

Beyond these phase-dependent dissociations, cognitive load systematically modulated the magnitude of striatal responses across task phases. The opposite scaling of activation across task phases appears to reflect how cognitive effort is distributed over time rather than a change in the roles of striosome- and matrix-like compartments. In the 0-back condition, the task is simple and the cue clearly signals what response will be required. As a result, the cue itself carries substantial information and evokes strong preparatory responses. In contrast, during the 2-back condition maintenance of the presented stimulus is a persistent task demand. Since activation is continuously present and less dependent on the brief cue block, the cue may encode less information, leading to reduced cue-related activation. Concurrently, activation during task execution increased with higher load, consistent with the greater demands of maintaining information, selecting responses, and monitoring performance during the 2-back task. As tasks become harder, the brain shifts effort from cue-based preparation to sustained execution, while preserving the distinct roles of the striosome- and matrix-like compartments.

When directly contrasting 2-back with 0-back conditions, both compartments demonstrated load-dependent increases in activation, but the effect was substantially stronger in matrix-like voxels (Figure 4). Matrix activation rose sharply with task difficulty across all stimulus categories, underscoring its close association with processes that support ongoing performance. Although prior anatomical studies have proposed roles for the matrix in encoding, manipulation, and execution of processes based on its extensive associative and sensorimotor connectivity (Flaherty and Graybiel 1993), these functions have not been directly demonstrated in humans or animals. Striosome-like activation also increased with load, though to a smaller degree, which may reflect contributions to motivational or evaluative aspects of performance under heightened cognitive challenge (Friedman, Homma et al. 2017, Amemori, Graybiel and Amemori 2021). Together, these results highlight a complementary division of labor: striosome-like voxels appear specialized for anticipatory and evaluative roles, while matrix-like voxels dominate the sustained processing required for ongoing and/or demanding memory and motor tasks (Okunomiya, Watanabe et al. 2025, Sadiq and Waugh 2025).

We next examined how compartmental activation differed between correct and error trials. Both striosome- and matrix-like voxels showed increased activation during correct 2-back trials relative to 0-back, indicating that both compartments scale similarly with task difficulty. During correct performance, matrix-like activation remained higher overall, consistent with its strong engagement during task execution. However, the key difference emerged during error trials: while both compartments showed reduced activation in the 2-back error condition, the decline was proportionally much larger in striosome-like voxels (71% decrease) than in matrix-like voxel (29% decrease). This pattern suggests that striosomal activation is particularly sensitive to trial accuracy, whereas matrix activation primarily reflects general task engagement or motor execution demands. Taken together, these findings highlight a functional dissociation: matrix activity scales with cognitive load, while striosome activity tracks performance accuracy, potentially indexing evaluative-or vigilance-related processes necessary for maintaining correct responses.

We also examined whether compartment-specific responses were driven by particular stimulus categories. In the 0-back condition, category-selectivity was most apparent in matrix-like voxels, which showed stronger activation for *body* part and *face* stimuli, while striosome-like activity remained relatively undifferentiated. This suggests that under low working memory demand, the matrix may be more sensitive to content-specific inputs that guide immediate responses. In contrast, under the 2-back condition, matrix activity became uniformly elevated across categories, consistent with its role in supporting sustained, domain-general task execution. Striosome-like voxels, however, displayed category-selective modulation at higher load, with significant enhancements for *place* stimuli and reduced activation for *body* part stimuli. These results indicate that the contributions of striosome-like voxels may become increasingly specialized under cognitive challenge, reflecting sensitivity to motivationally or contextually salient categories, while matrix responses shift toward generalized support of task performance.

In conclusion, this study provides the first evidence that in humans, the striatal compartments have specialized roles in working memory that vary with effort and the type of stimulus. Whereas our prior resting-state analyses demonstrated that these compartment-like voxels participate in segregated intrinsic functional networks (Sadiq, Funk, and Waugh, 2025), the current findings extend this framework by showing that compartmental organization also shapes activation dynamics under varying cognitive demands. Specifically, striosome-like voxels were preferentially engaged during cue-related processes, while matrix-like voxels dominated during sustained task execution and scaled robustly with increasing working memory load. Notably, this cue–execution dissociation parallels our prior motor task findings (Sadiq and Waugh 2025) suggesting that compartmental specialization may reflect a general organizing principle of striatal function across both motor and cognitive domains. These results support the emerging view that striatal microarchitecture contributes to phase-specific and demand-dependent specialization, with striosome- and matrix-like voxels differentially engaged across anticipation, execution, and performance accuracy. This compartmental specialization aligns with animal studies demonstrating that striosome and matrix circuits mediate distinct phases of motivated behavior. For instance, Friedman et al. (2015) showed that corticostriatal projections to striosomes are selectively recruited during decision-making under conflict, whereas matrix circuits support ongoing action execution (Friedman, Homma et al. 2015). Future studies will be necessary to link these patterns of functional activation to underlying cellular and network mechanisms, and to examine their generalizability across additional cognitive domains and in populations with neuropsychiatric diseases.

Our study combines structural connectivity–based parcellation, temporal decomposition of task epochs, and activation profiling to extend the functional characterization of striatal compartments beyond resting-state connectivity into real-time functional activation during working memory performance. This framework advances our understanding of how striosome- and matrix-specific networks support distinct components of cognitive control, including anticipatory processes, sustained information maintenance, and load-dependent execution. Importantly, these cognitive findings parallel our prior motor task fMRI results, suggesting that the compartmental division of labor represents a generalizable organizing principle across both motor and cognitive domains. These results further suggest that compartment-specific disruption may contribute to the cognitive and motor deficits observed in neuropsychiatric disorders involving the striatum and its associated networks.

## Data availability statement

Publicly available datasets were analyzed in this study. HCP data can be found here: https://www.humanconnectome.org/study/hcp-young-adult/document/1200-subjects-data-release. This dataset is BIDS compliant. The code, bait, seed, and exclusion masks necessary to complete striatal parcellation can be accessed here: github.com/jeff-waugh/Striatal-Connectivity-based-Parcellation.

## Author Contributions

AS: data acquisition and analysis, initial manuscript drafting, and critical revision of the manuscript. JW: data acquisition, analysis, and interpretation, initial manuscript drafting, and critical manuscript revision. All authors contributed to the article and approved the final version for submission.

